# A Cell Size-Dependent Competition Between Geometry and Polarity Governs Nuclear and Spindle positioning in Early Embryos

**DOI:** 10.64898/2026.01.23.701263

**Authors:** Aude Nommick, Macy Baboch, Celia Municio-Diaz, Jeremy Sallé, Remi Le Borgne, Nicolas Minc

## Abstract

Nuclei and mitotic spindles are actively positioned at defined locations within cells to regulate cell polarity, division and multicellular morphogenesis^1–4^. Forces generated by cytoskeleton networks regulate the positioning of these organelles and are commonly influenced by extrinsic cues such as cell geometry or polarity^5–12^. To date, however, most studies have investigated this problem in one given cell type, hampering our understanding for how mechanical systems that position nuclei and spindles may scale during multicellular development. We tracked the spatiotemporal behaviour of centrosomes, nuclei and spindles in early sea urchin embryos from the 1-cell to the ∼1000 cells blastula stage. We found that they are initially located at cell centers, but that they undergo a progressive decentration towards the embryo apical surface, as cells become smaller during development. This apical shift is mediated by microtubule (MTs) pulling forces which are influenced by both cell shapes and apical polarity domains. Using 3D mathematical models and embryo dissections, we propose that MT centering forces that derive from cell geometry decay in strength during development as a consequence of cell size reduction, allowing apical polarity decentering forces to take over. Our results support a self-organized scenario in which polarity cues progressively outcompete cell geometry, to modulate the overall balance of MT forces and pattern nuclear and spindle positioning throughout early embryo development.

## RESULTS AND DISCUSSION

### Nuclei and Mitotic Spindles become decentred during development

Early embryogenesis is characterized by a series of reductive cell divisions that are essential to establish the foundational architecture of developing organisms. These division patterns are regulated by the position and orientation of interphase asters organized around nuclei, and are often readjusted in mitosis through additional spindle translational or rotational motion ^9,10,13^. To characterize how the position of interphase asters and spindles evolve throughout early development, we performed sequential immunostaining, to visualize DNA, microtubules (MTs) and F-actin at different developmental stages of sea urchin *Paracentrotus lividus* embryos, from the 1-cell zygote stage to the ∼1000-cell blastula stage. Interestingly, this revealed that interphase asters bound to nuclei, as well as spindles, are located close to the cell center at early stages (2-cell to 4-cell), but that they progressively decenter towards the apical surface of the embryo at subsequent stages (8-cell to ∼1000-cell blastula stage). In addition, interphase aster pairs, as well as spindles appeared to align along an axis mostly parallel to the embryo surface, indicating that divisions are oriented in a planar manner throughout early development (Fig 1A-B). We confirmed these observations using live imaging of embryos injected with an NLS-GFP construct to mark nuclei, or with mRNA encoding for TMBD-mStayGold and LifeAct-mScarlett3 to image MTs and F-actin respectively, and visualize asters, spindles and cell shapes (Fig S1A-C and Movie S1).

**Figure 1.**
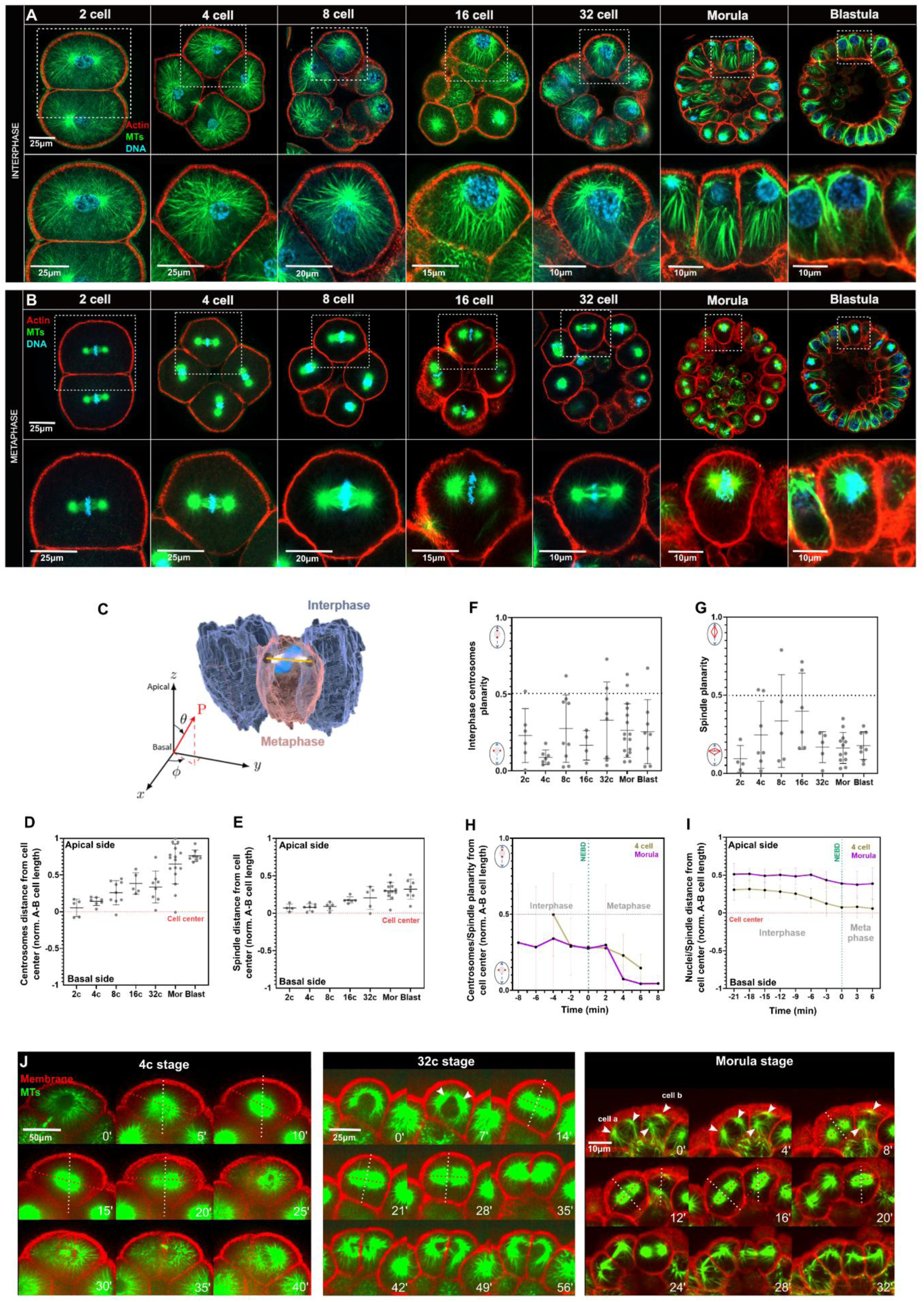
Patterns of nuclear and spindle positioning during early embryo development. **(A-B)** Confocal single-slice acquisitions of fixed sea urchin embryos at successive developmental stages in interphase (A) and metaphase (B). Embryos were stained for F-actin (red), Microtubules (green) and DNA (blue). Insets highlight individual blastomeres. **(C)** Three-dimensional segmentation of three blastomeres (blue = interphase; red = metaphase). The vector P (in red) represents the orientation of the mitotic spindle axis in a three-dimensional reference frame associated with the apico-basal axis of the cell. **(D-E)** Quantification of nuclear (C) and spindle (D) distances from the cell center (red dotted line), normalized to the apico-basal cell length across developmental stages. (n= 4-10 cells/stage). **(F-G)** Quantification of interphase centrosome-to-centrosome pairs (F) and spindle (G) orientation. Values approaching 1 indicate an orientation parallel to the apical–basal axis, and values close to 0 indicate a planar orientation orthogonal to the AB axis. **(H)** Time evolution of centrosome pair orientation from interphase to metaphase at 4-cell or morula stages (n=4 cells/stage). **(I)** Time evolution of nuclei and spindle positions from interphase to metaphase at 4-cell and morula stages (n=4 cells/stage). **(J)** Time-lapse confocal projections of embryos injected with mRNA to express TMBD-mStayGold (MTs, green) and membrane-mCherry (Membrane, red). Left panel: transition from 4- to 8-cell stage, middle panel: from 32-cell to morula and right panel: from morula-to-morula stage. The red dashed line indicates spindle orientation and the white dashed line the apico-basal cell axis. White arrows point to the position of centrosomes in interphase/prophase. Scale bar lengths are indicated in corresponding panels. Error bars represent +/- S.D.

In order to quantify the position and orientation of interphase aster pairs and mitotic spindles, we segmented images from fixed embryos at all stages in 3D (Fig 1C). This showed that interphase centrosomes at aster centers, progressively shift towards the apical surface to reach a mean decentering distance at the blastula stage of ∼75% of the apico-basal cell radius (Fig 1D and Fig S1D). Similar decentration dynamics was obtained for interphase nuclei, but with decentering distances shorter than for centrosomes, reflecting the more apical location of interphase centrosomes as compared to nuclei centers (Fig 1A, S1D-E). This analysis also confirmed a progressive decentration of spindles, but these were located closer to the cell center, as compared to interphase asters, reaching a decentering distance of only ∼32% of the apico-basal cell radius by the blastula stage (Fig 1E). Accordingly, live imaging of embryos injected with mRNA encoding for TMBD-mStayGold and Membrane-mCherry, confirmed a systematic recentering dynamics of centrosomes between interphase and metaphase (Fig 1I-J).

We also used this 3D analysis in fixed cells to compute the dot product between interphase centrosomes or spindle axis with the apico-basal (AB) axis. Dot products were generally below 0.3-0.4 corresponding to angles above ∼60-70°, confirming that centrosome axes lie mostly orthogonal to the AB axis, in the plane of the embryo surface. Dot products were generally smaller in metaphase, especially at late stages, suggesting slight realignments of centrosome/spindle axis in metaphase, that we also confirmed by live imaging (Fig 1F-H, 1J and Fig S1F). All together these results show that interphase asters, nuclei and mitotic spindles follow a stereotyped positioning dynamics during early development, marked by a continuous planar alignment and a progressive apical decentration.

### Microtubules promote the apical decentration of nuclei

To understand how centrosomes progressively decenter to the apical side of the embryo, we characterized cytoskeleton organization at early and late stages of development. In interphase, astral MTs filled the whole cell volume at all stages, indicating they may function to probe cell geometry and/or cortical polarity^9,14,15^. In metaphase, astral MTs were significantly shorter, and appeared to be mostly uncoupled form the cell surface at early stages, reaching the cortex only at stages beyond the 8- or 16- cell stage^16^ (Fig 1A and 1B). F-actin mostly decorated the cortex, but we noted the formation of short actin cables emanating specifically from the apical cortex. These cables appeared to prolong microvilli at the apical surface of cells, and were generally longer in interphase as compared to metaphase (Fig 2A and Fig S2A). Importantly, these cables were readily observed as early as in the 2-cell stage, reflecting the existence of an internal apico-basal polarity even at early developmental stages, as previously proposed^17,18^ (Fig 2A). In addition to F-actin, we also imaged phospho-MyoII, and observed that it accumulated at the apical cortex in interphase and metaphase at all stages^19^. This indicates that apical cortices may exhibit higher actomyosin contractile activity, as compared to basal ones, at all developmental stages (Fig 2B). Finally, we also imaged organelle content in the cytoplasm with transmission electron microscopy, and found no evidence for organelle accumulation at the basal vs apical pole; indicating that cytoplasm content in organelles may not bias centrosome or nuclear positioning, as has been proposed for other embryos^13,14^ (Fig S2B-C).

**Figure 2.**
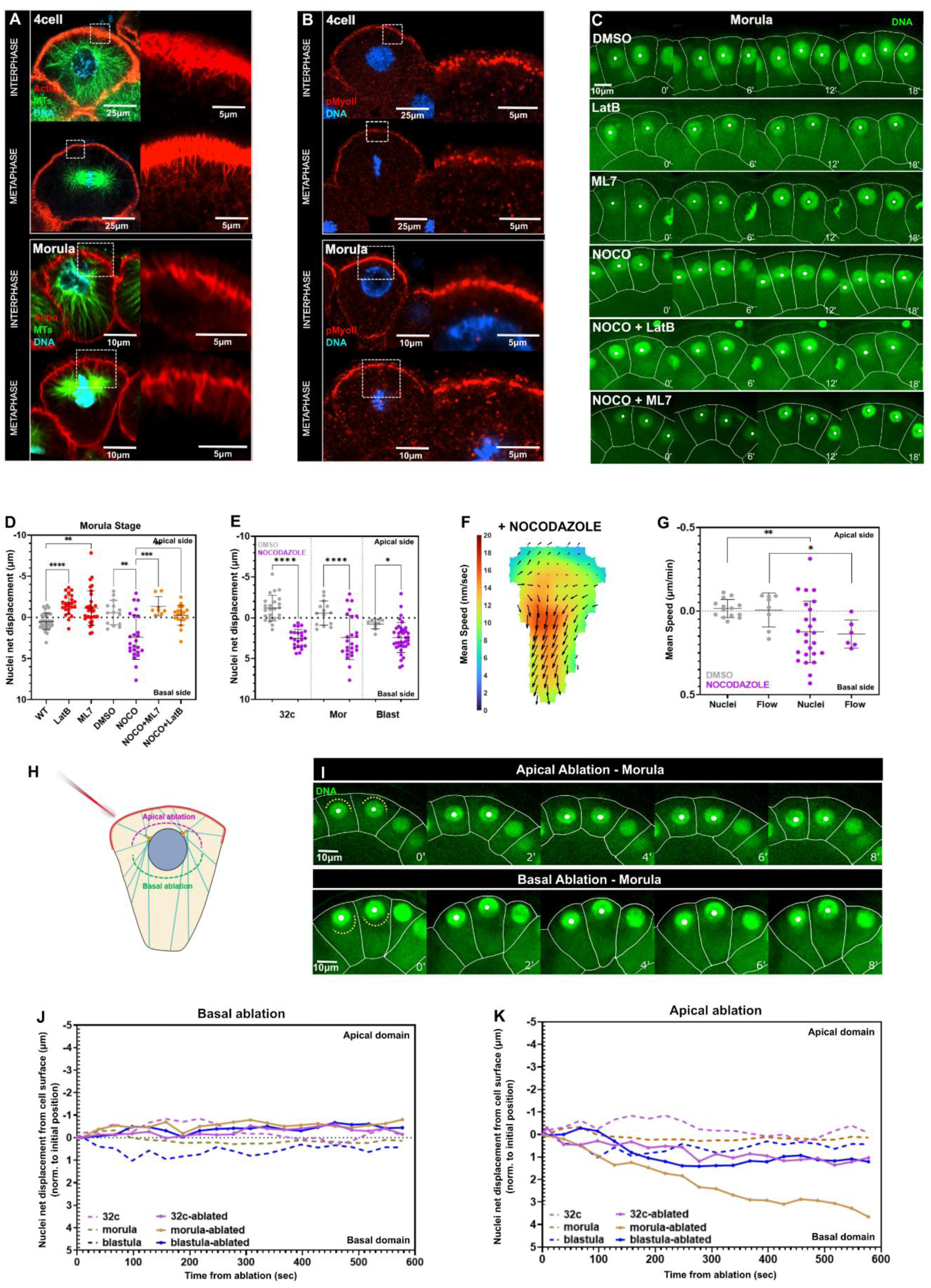
Microtubule pulling forces regulate centrosome apical decentration at late stages. **(A)** Confocal single slices of fixed sea urchin blastomeres at 4-cell and morula stages, in interphase and metaphase. Embryos were stained for F-actin (red), MTs (green), and DNA (blue). Insets highlight actin cables growing from the apical cell domain. **(B)** Confocal single slices of fixed sea urchin blastomeres at 4-cell and morula stages, in interphase and metaphase. Embryos were stained for Phospho-MyoII (red), and DNA (blue). Insets highlight apical domains enriched in phospho-myosin. **(C)** Confocal time-lapse single slice of morula-stage embryos incubated in Hoechst to label DNA (green) and treated with the indicated chemicals. White lines delineate cell membranes, and white spots mark nuclei centers. **(D)** Quantification of nuclear net displacement after 15-30 minutes (from nuclei appearance until the end of interphase) in embryos treated with indicated chemicals (n > 10 cells/condition). **(E)** Quantification of nuclear net displacement after 15-30 minutes (from nuclei appearance until the end of interphase) in embryos treated with nocodazole or DMSO at the indicated developmental stages indicated chemicals (n > 10 cells/stage). **(F)** Cytoplasm flows, analysed by tracking yolk granules in confocal imaging, within a single cell in an embryo treated with nocodazole at the morula stage. Vectors represent flow direction and amplitude. **(G)** Nuclear and cytoplasm flow speeds in the indicated condition in embryos at the morula sage (n > 10 cells for nuclei speeds, n= 6-8 cells for flow speeds). **(H)** Scheme of a cell at morula stage depicting regions in which MTs were ablated with a laser beam. **(I)** Time-lapse confocal single slice of morula-stage embryos incubated in Hoechst to label DNA (green) after MT laser ablation. White lines delineate cell membranes, yellow dashed lines indicate ablation regions and white spots mark nuclei centers. (J-K) Nuclear displacements after apical or basal MT laser ablation. Time 0 correspond to the moment of ablation. The curves represent the mean of all nuclei displacements (dashed lines: control non-ablated, solid lines: ablated). (n > 10 cells/condition). Scale bar lengths are indicated in corresponding panels. Error bars represent +/- S.D. Results were compared using a two-tailed Mann–Whitney test. P-values are indicated as *, P < 0.05, **, P < 0.01, ***, P < 0.001, ****, P < 0.0001.

The decentred apical location of centrosomes suggests the dominance of an apically-directed force. To identify which cytoskeletal element may generate this asymmetric force, we treated late-stage embryos with low doses of specific cytoskeleton inhibitors. For this, we labelled nuclei with Hoechst, and tracked nuclear movements, as a proxy for aster positioning, with live imaging following treatments with inhibitors. Depolymerizing F-actin with a low dose of Latrunculin B, or affecting myosin activity with an ML7 inhibitor led to a net apically-directed motion of nuclei. This suggests that the apical cortex enriched in actomyosin, may contract to exert a basally-directed force that shifts nuclei away from the apical cortex^20,21^. In sharp contrast, treating cells with low doses of nocodazole to depolymerize MTs led to a rapid basally-directed motion of nuclei, indicating that MTs are the primary force generators to decenter nuclei and centrosomes towards the apex. Nocodazole-induced nuclei movements were abolished by adding Latrunculin B or ML7 to inhibit actomyosin activity, suggesting that basally-directed movements of nuclei in the absence of MTs results from actomyosin contractile activity at the apex (Fig 2C-E, Fig S2D and Movie S2). Accordingly, in nocodazole treated embryos we observed cytoplasm flows directed towards the basal poles, presumably deriving from apical actomyosin contraction, with speeds similar to that of descending nuclei (Fig 2F-G)^20,21^. Together, these results suggest a predominant role for the MT cytoskeleton in decentering centrosomes towards the apex, with some minor basally directed opposed force exerted by actomyosin contractility.

We next asked if MTs may exert pushing or pulling forces to decenter nuclei and centrosomes towards the apical pole. In general, astral MTs appeared relatively straight in interphase asters, suggesting they may be subjected to tensile forces (Fig 1A and Fig S1A). To more directly assay force direction, we performed laser ablation of MTs. Ablating MTs on the apical side, led to a basally-directed movement of nuclei at all stages tested. Conversely, MT ablation from the basal side led to an apically-directed movement of nuclei, albeit smaller in amplitude as compared to basal ablations, presumably because nuclear motion may be limited by the apical cortex (Fig 2H–2K). Therefore, these results indicate that MTs primarily exert pulling forces, that may vary in amplitude and/or asymmetry over development to progressively shift centrosome positions towards the surface of the embryo.

### Cell geometry is not sufficient to account for centrosome position and orientation throughout embryo development

Cell geometry has been shown to serve as an important cue to position centrosomes in the cell center and orient them along the long cell shape axis, especially in the context of early embryo development ^9,13,14,22^. To explore the role of cell shape, we quantified cellular aspect ratios, defined as the apico-basal cell length divided by the transverse length. This revealed that cells evolve from a flattened shape larger along the transverse axis at early stages, to a more rod-like shape now elongated along the AB axis at later stages (Fig 3A). To test if these geometrical changes could account for modulations in centrosome positioning and orientation observed during development, we ran 3D simulations. Using 3D segmentations of individual cell contours as input for cell geometries, we placed centrosome pairs at random positions and orientations and used a gradient descent simulation, that compute force and torque exert generated by astral MTs on centrosomes to identify mechanical equilibria^14,23^. This first model was based on pure geometrical considerations by assuming that astral MTs fill and probe the whole cell volume, by exerting pulling forces scaled to the cube of their length ^9,14,24,25^. In metaphase, these simulations revealed that geometry can serve as a good predictor of spindle position and orientation at early 2-cell or 4-cell stages, but that these geometrical predictions depart from experimental results at later stages, with spindles largely deviating from the cell geometrical center and also aligning along the shortest cell shape axis, in sharp contradiction to model predictions (Fig 3B-D).

**Figure 3.**
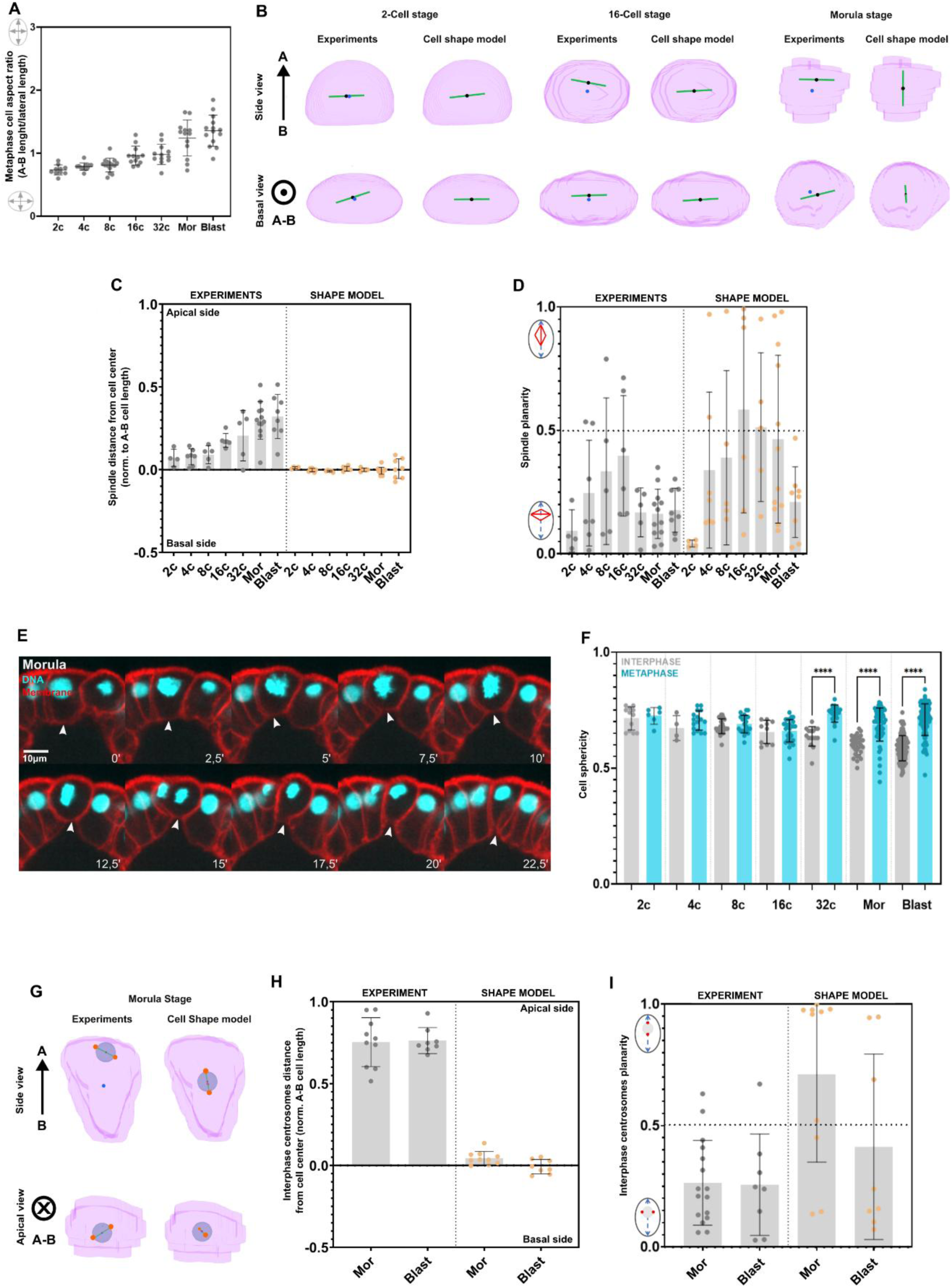
Cell geometry is not sufficient to explain centrosome positioning during early development. **(A)** Quantification of metaphase cell aspect ratio defined as the apico-basal cell length divided by lateral cell length (n>10 cells/condition). **(B)** 3D model predictions of metaphase spindle orientation and position at different developmental stages. The green line marks the spindle axis, the blue dot, the cell center of mass and the black dot the spindle center. **(C-D)** Experimental results (grey dots) and 3D model predictions for the same cells (orange dots) of spindle distance from the cell center normalized to apico-basal cell length (C) and spindle planarity (D) at different developmental stages**. (E)** Confocal time-lapse projections of morula-stage embryo expressing H2B (DNA, cyan) and membrane-mScarlet3 (red). Arrowheads indicate the basal cell domain that rounds up during mitosis. **(F)** Quantification of cell sphericity across different stages in interphase (grey) and metaphase (blue) (n=6-60 cells/stage). **(G)** 3D model predictions of interphase centrosome-pairs orientation and position. Orange dots mark centrosomes; the grey circle, the nucleus; the blue dot, the cell center of mass and the black dot the nucleus center. **(H)** Experimental results (grey dots) and 3D model predictions for the same cells of interphase centrosome-pairs position (H) and planarity (I) at morula and blastula stage (n=8-10 cells/stage). Scale bar lengths are indicated in corresponding panels. Error bars represent +/- S.D. Results were compared using a two-tailed Mann–Whitney test. P-values are indicated as ****, P < 0.0001.

In addition, we observed some level of shape changes between interphase and mitosis, with cells undergoing partial rounding at late stages beyond the 32-cell stage (Fig 3E-F). This prompted us to run simulations based on interphase cell shapes at these late stages. Here again, model prediction based on 3D interphase cell geometry failed to account for nuclear/centrosome apical locations and for their planar orientations (Fig 3G-I). We conclude that cell geometry may constitute an important element to control centrosome positioning at early developmental stages^9,13,14^, but that it is insufficient to explain centrosome decentration and planar orientation at later stages.

### A titrated competition between polarity and geometry predicts centrosome positioning throughout development

The progressive apical decentration of centrosomes suggested a relative increase in apically directed MT forces during early development, likely regulated by apical polarity domains^26–28^. As mentioned above, our observation of a markedly different actin cortical network on the apical pole, as well as the apical enrichment in phospho-MyoII, indicates the emergence of apico-basal polarity as early as in the 2-cell stage^17^. To more directly monitor upstream regulators of apico-basal polarity, we stained embryos at all stages for phospho-PKCζ (protein kinase C zeta). PKCζ is an atypical isoform of protein kinase C, conserved across species, that plays a central role in the establishment and maintenance of apico-basal polarity ^18,29–31^. This confirmed the presence of polarized membrane domains, as early as in the 2-cell stage, with a marked enrichment of the apical surface, that persisted throughout development. Apical enrichment in p-PKCζ as compared to the basal one, was constant around ∼2 fold throughout development, ruling out any major developmental maturation of AB polarity. Importantly, these p-PKCζ domains colocalized with apical zones enriched in actin cables, indicating that these apical protrusions indeed reflect the presence of upstream polarity effectors. Therefore, these results confirm the existence of *bona fide* AB polarity established from the beginning of embryo development.

In order to test the contribution of apical polar zones to centrosome positioning, we turned back to 3D simulations. We added to the 3D geometrical model a polar domain, with size and position inferred from 3D images of apical zones enriched in F-actin protrusions, and assumed that MTs contacting this domain exert an additional force as compared to other MTs in the cell (Fig 4D). As such, in the model, all MTs pull with a force that scales to the cube of their length to account for a geometrical effect based on probing cell volume, and MTs contacting the apical polar domain exert an additional force scaled to the square of MT length, to model a “surface-based” pulling ^14,32^. The ratio between these two contributions is embedded in the value of a single parameter, β. Using constant low or high values for β at different stages, led to poor predictions of the model, with an underestimation of centrosome apical decentration or a premature one, respectively (Fig S3A-C). However, volume-based MT pulling amplitude may decay with cell size, given that the length of MTs become shorter in smaller cells. As such, we inversely scaled β to cell volume, to account for volume reduction of blastomeres and consequent decrease in the ratio between geometrical-based centring forces and polarity based decentring forces during development (Fig 4D and 4F). Remarkably, without adjusting any parameter between simulations, this model reproduced the complete trend of progressive decentration of interphase centrosomes and spindles, as well as their planar orientation at all stages (Fig 4E, 4G-J). Therefore, these results suggest that a progressive competition between geometry and polarity, regulated by cell volume, may account for the full spatiotemporal pattern of centrosome positions during early development.

**Figure 4.**
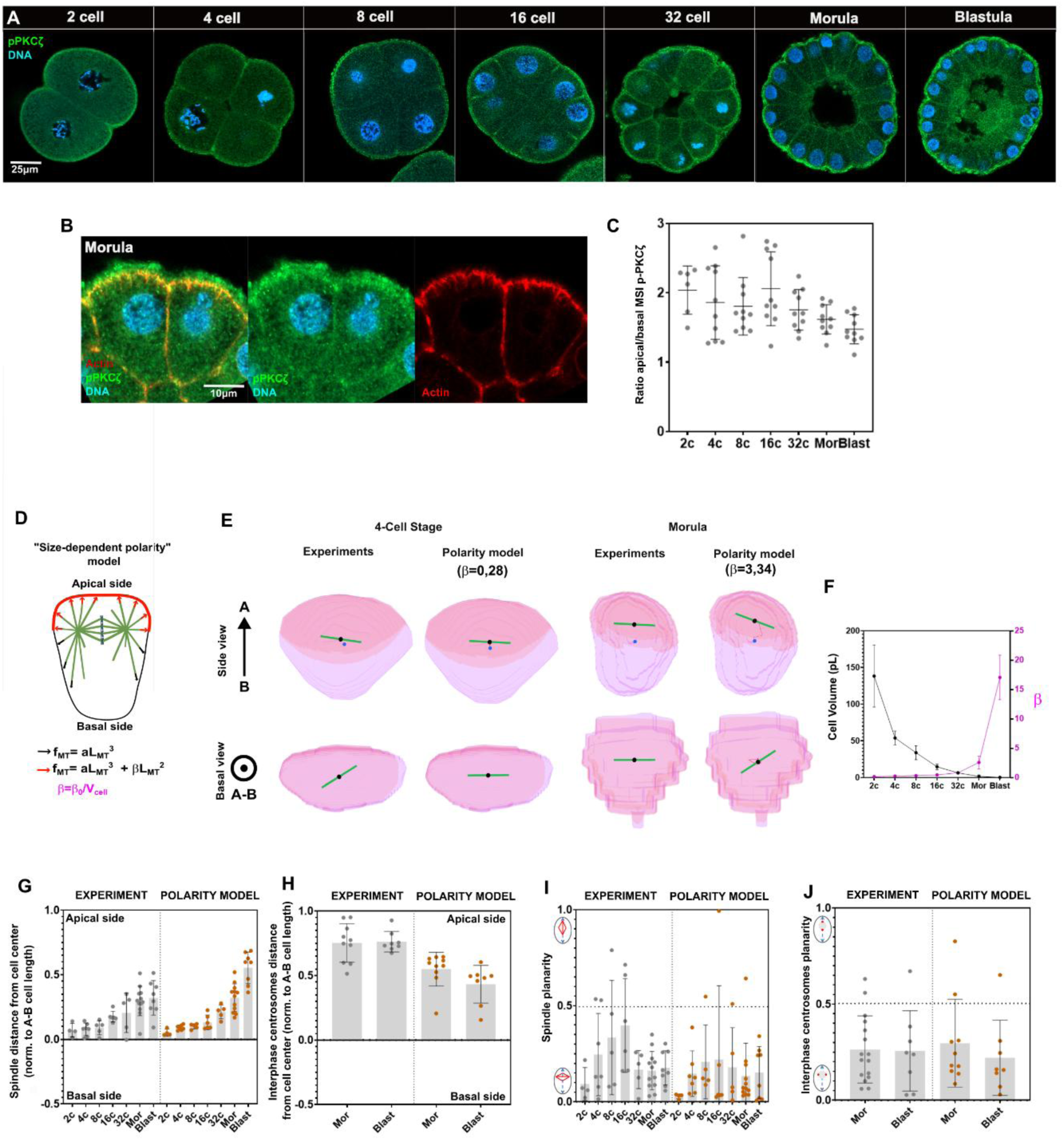
Size-dependent competition between apical polarity and cell geometry accounts for centrosome positioning dynamics during development. **(A)** Confocal projections of fixed embryos at successive developmental stages stained for the phosphorylated form of PKCζ (p-PKCζ, green) and Hoechst (DNA, blue). **(B)** Confocal projections of fixed blastomeres at the morula stage, stained for p-PKCζ (green), Phalloidin (F-actin, red) and Hoechst (DNA, blue). **(C)** Quantification of the ratio between apical and basal p-PKCζ mean signal intensity (MSI) (n=10 cells/stage). **(D)** Scheme of the size-dependent polarity model for centrosome positioning: All astral MTs pull with a force proportional to the cube of their length, f_MT_ =aL_MT_^3^, with a fixed constant, but MTs that contact the apical domain pull with an additional force so that f_MT_ =aL_MT_^3^ +β L_MT_^2^. To account for volume reduction of blastomeres during development, β is inversely scaled to cell volume. **(E)** 3D model predictions of metaphase spindle orientation and position at different stages. The green line marks the spindle axis, the blue dot, the cell center of mass and the black dot the spindle center, the pink area delineates the apical polar domain **(F)** Evolution of measured cell volume during development, and consequent impact on values of the parameter β as a function of developmental stages (n=4-10 cells/stage). **(G)** Experimental results (grey dots) and 3D model predictions for the same cells (orange dots) of spindle distance from the cell center normalized to apico-basal cell length (n= 4-10 cells/stage). **(H)** Experimental results (grey dots) and 3D model predictions for the same cells (orange dots) of interphase centrosome distance from the cell center normalized to apico-basal cell length, at morula and blastula stages (n=8-12 cells/stage). **(I)** Experimental results (grey dots) and 3D model predictions for the same cells (orange dots) of spindle planarity (n= 4-10 cells/stage). **(J)** Experimental results (grey dots) and 3D model predictions for the same cells (orange dots) of interphase centrosome pairs planarity (n=8-12 cells/stage). Scale bar lengths are indicated in corresponding panels. Error bars represent +/- S.D.

### Cell size is a key regulator of nuclear position

A direct prediction of the model, is that a smaller cell volume should yield to a more pronounced centrosome apical shift, at the same developmental stage. To test this prediction, we performed microdissections of unfertilized eggs, and fertilized these cut eggs to produce smaller embryos, with smaller cells and tracked nuclear positioning (Fig 5A-B). Quantification of cell volumes from immunostaining images, confirmed a marked reduction of ∼1.5 fold on average, in cut embryos as compared to control embryos from the same batch at the same developmental stage. Remarkably, this size reduction, caused a marked precocious apical shift in nuclear positioning, with nuclei at the 4- or 8- cell stage in cut embryos being as close to the apical cortex as those of 16- or 32-cell stages in control non-cut embryos (Fig 5D). We noted however, that this impact of cell size reduction saturated at later stages, presumably because interphase asters and nuclei reach a minimal distance to the apex set by the presence of the apical cortex (Fig 2D). Finally, by plotting nuclear apical shifts as a function of cell volumes for control and cut-embryos revealed a clear negative dose-dependence correlation (correlation coefficient; −0.71), strongly supporting the impact of cell size on MT mechanical force balance that position centrosomes and nuclei (Fig 5E).

**Figure 5.**
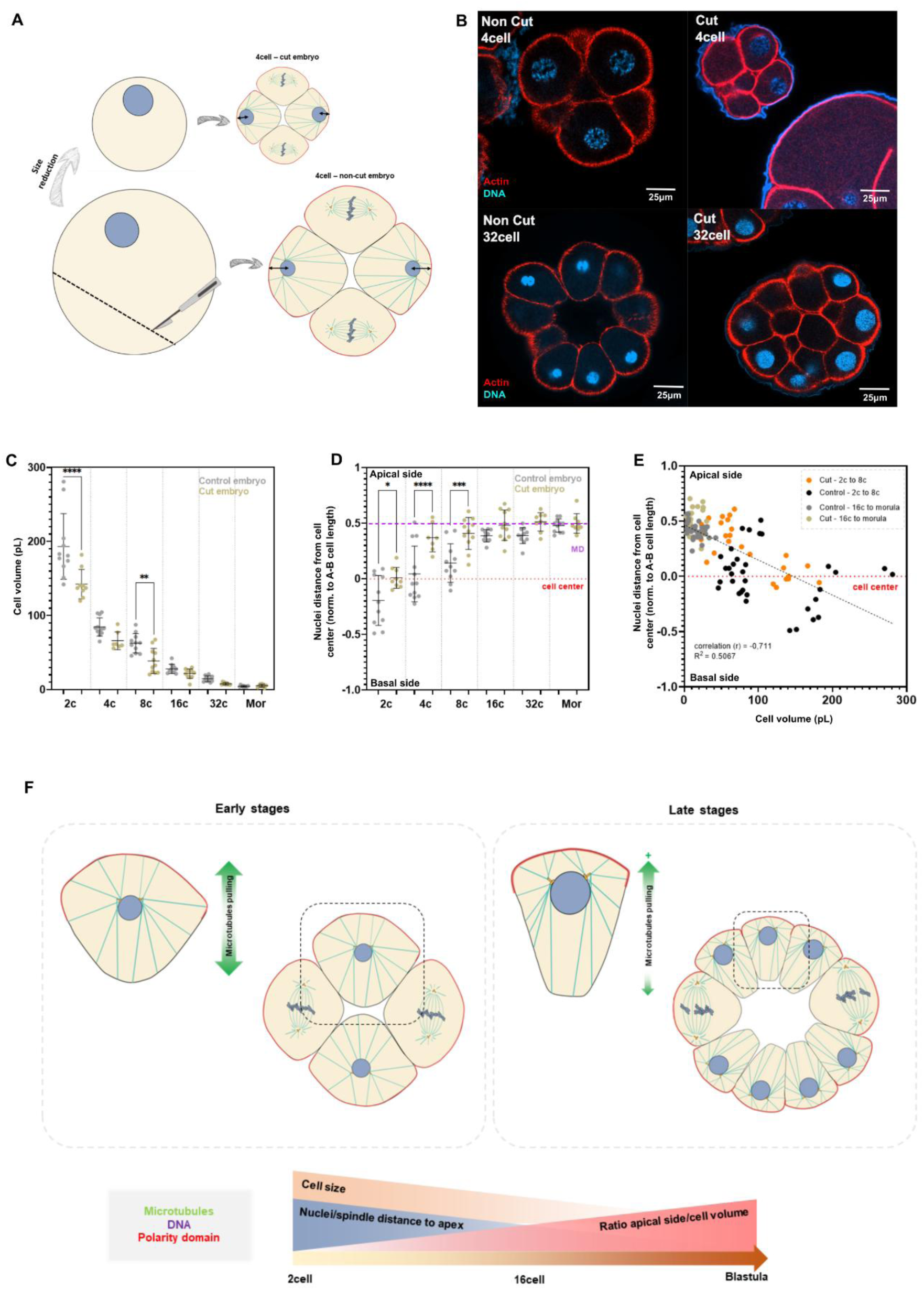
Cell size directly impacts nuclear positioning. **(A)** Scheme representing microdissection experiments in unfertilized sea urchin eggs. Eggs were mechanically cut with a micro-needle, and then fertilized. Nuclei position was tracked at different stages and compared to non-cut embryos from the same batch. **(B)** Confocal acquisitions of fixed intact (“non-cut”) and cut embryos at the 4- and 32-cell stages. Embryos were stained with Phalloidin (F-actin, red) and Hoechst (DNA, blue). **(C)** Comparison of blastomere volumes in cut vs non-cut embryos (n=3-6 embryos/stage). **(D)** Nuclear distance from cell center normalized to the apico-basal cell length on cut vs control embryos at different developmental stages (n=3-6 embryos/stage). **(E)** Nuclear distance from cell center normalized to the apico-basal cell length plotted as a function of cell volume for individual blastomeres in the indicated stages and conditions. The dotted line is a linear regression. **(F)** Proposed “scaling” model for centrosome positioning in early embryos: As cells reduce in volume, during development, the amplitude of MT centering forces that depend on MT length reduces, rendering apical surface decentering forces to take over and progressively bring centrosomes, nuclei and spindles closer to the embryo surface.

The mechanical elements that position and orient centrosomes in interphase or metaphase, are fundamental regulators of embryo development and tissue morphogenesis, and have been investigated in numerous single-cell types ^7–9,33^. Here, we systematically mapped centrosome, nuclear and spindle positions throughout the course of multicellular embryo development. We found that centrosomes follow stereotypical graded positional mechanisms, with a progressive apical shift and a near constant planar orientation through early development. Our results suggest that a single rule based on the competition between the influence of AB polarity domains and cell shape on astral MT forces, titrated by cell volume reduction, is sufficient to account for these patterns. Therefore, an important output of this study is to put forward a “cell size scaling” model for MT force balance and consequent nuclear and spindle positioning. Size scaling is fundamental for early embryo development^34,35^. It ensures that organelles such as nuclei or spindles will assume adequate dimensions to fulfil cellular functions, or that dynamic processes such as cytokinesis or chromosome segregation are executed within the same timing through embryo development ^36–41^. We propose that size scaling of MT forces for centrosome positioning may be essential to ensure that spindles end up apically located in forming columnar morula/blastula cells, to promote faithful bipolar spindle formation, chromosome segregation or monolayered blastoderm tissue architecture^23,42^. Future work addressing in quantitative terms how cytoskeletal forces intersects with cell size will bring important insights into mechanisms of organelle positioning and tissue morphogenesis.

## RESOURCE AVAILABILITY

### Lead contact

Further information and requests for resources and reagents should be directed to and will be fulfilled by the lead contact, Nicolas Minc.

### Materials availability

This study did not generate new unique reagents.

### Data and code availability

- All data needed to evaluate the conclusions in the paper are present in the paper and/or the supplemental information.
- This paper does not report any original code.
- Any additional information required to reanalyse the data reported in this work is available from the lead contact upon request.

## ACKNOWLEDGMENTS

We thank all members of the Minc team for discussion and technical help. We thank J. Chenevert (LBVD, Villefranche); A. Mc Dougall (LBVD, Villefranche); V. Barone (Stanford U.) and D. Levy (U. Wyoming) for sharing reagents. A.N. was supported by a post-doc fellowship from the ARC foundation (PDF2021070004082). We acknowledge the ImagoSeine core facility of the Institut Jacques Monod, member of the France BioImaging infrastructure (https://ror.org/01y7vt929) supported by the French National Research Agency (ANR-24-INBS-0005 FBI BIOGEN) and GIS-IBiSA. This work was supported by the Centre National de la Recherche Scientifique (CNRS), the Université Paris Cité, and grants from La Ligue Contre le Cancer (EL2021.LNCC/ NiM), the Agence Nationale pour la Recherche (ANR, “TiMecaDev”) and the Fondation Bettencourt Schueller (“Impulscience”), to N.M.

## AUTHOR CONTRIBUTIONS

Conceptualization, N.M., and A.N.; Methodology, N.M., R. L.B., C. M-D., J.S., M.B, A.N.,

Writing –Original Draft, N.M., and A.N Draft Editing. N.M., R. L.B., C. M-D., J.S., M.B, A.N.

## STAR METHODS

## Experimental model and subject details

### Sea urchin gametes

Purple sea urchins (*Paracentrotus lividus*) were obtained from Roscoff Marine station (France) and maintained at 16°C in aquariums of artificial sea water (ASW; Reef Crystals, Instant Ocean). Female and male gametes were collected by opening animals, or by intracoelomic injection of 2mL 0.5 M KCl. Eggs were collected and washed in ASW.F (artificial sea water filtered). Sperm was collected with similar methods, and kept at 4°C and used within 2-4 days after collection. When eggs were injected, they were first filtered 3 times through a 70-μm Nitex (VWR) mesh to remove the vitelline membrane and facilitate adhesion to the protamine-coated dish. Unfertilized eggs were incubated at 16°C in ASW.F until use the same day of collection. If the final purpose was to fix embryos for immunostaining, they were fertilized in ASW.F + PABA 0.274mg/mL for 2 min to remove the fertilization membrane and facilitate antibody (Ab) penetrance. Then, they were filtered through an 80-μm Nitex (VWR), and incubated in ASW.F at 16°C to develop until the desired stage prior to fixation.

## Method details

### Immunostaining

Embryos were fixed either directly on protamine-coated dishes or in bulk in falcon tubes in FA buffer (100mM HEPES-pH 6.9, 50 mM EGTA, 10mM MgSo4, 2% Formaldehyde, 0.2% Glutaraldehyde, 0.2% Triton X-100, 400mM Dextrose) for 70min at room temperature (RT). Then, they were washed 3X in PBT and stored at 4°C or used directly for immunostaining. First, to remove auto-fluorescence, they were incubated 30 min in PBS 1x + 0.1%NaBH4 and washed 2X in PBS for 10min. Then, embryos were blocked in PBT + 0.1%BSA + 5% Goat Serum for 30min at RT. The blocking solution was replaced by the primary antibody (Ab) diluted in PBT + BSA 0.1% and embryos were incubated ON at 4C°. The next day, they were washed 3X in PBT for 30min and incubated in the solution containing the secondary Ab, and Hoechst (1/1000) to label DNA diluted in PBT for 3h at RT in the dark. Finally, they were washed 1X in PBT 30min, 1X in PBS and finally in PBS 1X + 50% Glycerol and let to sediment for 30 min until mounting. Embryos were mounted on a glass dish (50mm Dish, No. 1.5 coverslip, 14mm Glass DiaMTser, MatTek Corporation©) in glycerol-based Mounting Medium, and covered with a 18×18mm coverslip sealed with nail polish. All antibodies used and their concentrations are listed in the Key Resource table.

### Protein synthesis

Recombinant GST-GFP-NLS and GST-mCherry-NLS were expressed and purified as previously described ^38,43^. Protein stock concentrations were generally around ∼1-10 mg/ml.

### mRNA synthesis

Capped-mRNA were synthetized with the mMessage mMachine kit (ThermoFisher) and purified with the MEGAClearPURIF kit (ThermoFisher). To increase mRNA stability and translation efficiency a Poly(A) sequence was added with the Poly(A) Tailing KIT (Ambion).

All constructs are listed in the Key Resource table.

### Egg injection

Injections experiments were performed on an inverted epifluorescence microscope (TI-Eclipse, Nikon) combined with a complementary metal–oxide–semiconductor (CMOS) camera (Hamamatsu), using 20x and 10x dry objectives (Apo, NA 0.75, Nikon). The microscope is operated with Micro-Manager (Open Imaging), at a stabilized room temperature (18–20°C).

Injections were done with capillaries TW100F-4 (borosilicate glass capillaries 1mm diameter) (World Precision Instruments, WPI), previously pulled (P-1000, Sutter Instrument) and tip-ground with the MicroGrinder (EG-402 Narishige) at an angle of 40°. Eggs were adhered to glass dishes (µ-Dish 35mm 1.5 low, IBIDI) coated with 1% protamine and placed on the inverted epifluorescent microscope, equipped with a micromanipulator (Injectman 4, Eppendorf) and a micro-injection system (FeMTsoJet 4, Eppendorf). After injections, eggs were fertilized with 1 drop of diluted sperm, and imaged for *in vivo* experiments, fixed, or incubated at 16°-20°C until the desired stage.

Before each experiment, the −80°C RNA stock aliquot was centrifuged 5min at 19000rpm at 4°C/RT. Then, it was dissolved at the desired concentration in RNAse-free-H2O.

### Chemical inhibitors

The concentrations and incubation times used for different cytoskeleton inhibitors were: Nocodazole 20 µM for 5min, ML7 100 µM for 15min, Latrunculin B 0.5 µM for 15min. Drugs were diluted to their final concentration from a 100X stock in DMSO. For immunostainings, embryos were developed in ASW.F, and transferred in a tube containing 5mL ASW.F supplemented with inhibitors, and let to incubate for varying time depending on the inhibitor, prior to fixation. For live imaging, embryos were developed on a dish in ASW.F until the desired stage. Then, the drug was diluted in 500µl of ASW.F and added to the dish. All drug concentrations and incubation times were calibrated in order to minimize toxicity and embryo mortality. The control conditions used to compare drug effects have been ASW.F or ASW.F + DMSO, depending on the medium in which the original drug stock was dissolved.

### Egg bisection

Non-fertilized eggs were adhered to glass dishes (µ-Dish 35mm 1.5 low, IBIDI) coated with 1% protamine and incubated in ASW.F + Hoechst. A micro dissecting knife (Item No. 10055-12 FST) was used for cutting the embryos under a binocular microscope (Nikon SMZ1270). Cut eggs were then fertilized by adding one drop of diluted sperm and incubated at 16°C until the desired stage of imaging or fixation.

### Imaging

Live-imaging and immunostaining of fixed embryos were performed on three different microscopes: (1) A spinning-disk confocal microscope (TI-Eclipse, Nikon) equipped with a Yokogawa CSU-X1FW spinning head, and an EM-CCD camera (Photometrics, PRISM BSI) and a 40xWater, 40xOil or 60xOil objectives; (2) A spinning-disk confocal microscope (TI-Eclipse, Nikon) equipped with a Yokogawa CSU-W1 spinning head, and an EM-CCD camera (Photometrics, KINETIX 22) and a 60xWater objective; (3) A confocal microscope Zeiss LSM980, coupled to the Airyscan 2 module equipped with 63xOil and 40xWater objectives. Imaging was performed at room temperature (between 18-25°C).

### Electron Microscopy

Sea urchin embryos, at the morula stage, were processed by high-pressure freezing and freeze substitution. The samples were placed in a 200 µm deep flat specimen carrier, 1.5mm dia. The sample was immediately high-pressure frozen (EMPact2, Leica). Frozen carriers were transferred to a cryotube containing 1 mL of cryo-substitution mixture, consisting of 2% osmium, 0.1% glutaraldehyde (GA) and 0.1 % uranyl acetate (UA) in acetone. Since GA and UA were diluted from an aqueous solution, the final mix included 2.9 % H2O. Freeze-substitution was performed in an AFS2 (Leica), at −90 °C for 50 hours, then the temperature was gradually increased to −30°C, this temperature was maintained for 24 hours. Finally, the temperature was gradually raised to +15 °C. The freeze-substitution mixture was removed and specimens were washed in two 15 min acetone baths and infiltrated with epoxy resin (Agar, low viscosity resin) in baths of increasing concentration of resin, up to 100%. Polymerization was done at 60°C for 24h. 70-nm-ultrathin sections were cut using an EM UC6 ultramicrotome (Leica) and collected on slot grids. The sections were post-stained in 2% aqueous uranyl acetate and lead citrate (Reynold’s solution) and observed at 120 kV with a Tecnai12 transmission electron microscope (ThermoFisher Scientific) equipped with a 4K OneView camera (Gatan)

### Laser Ablation

Single Z-plane laser ablation experiments were performed using a 355 nm UV laser integrated into a spinning disk confocal microscope (iLas Pulse system, MetaMorph software). Laser calibration was carried out every 30 minutes to correct for laser alignment drift over time and temperature. Embryos were adhered on glass dishes (µ-Dish 35mm 1.5 low, IBIDI) coated with 1% protamine with ASW.F + Hoechst under a 40XOil objective. Laser parameters were optimized for the experiment: 10 repetitions, 100% laser power, thickness 2. Time-lapse acquisitions were done every 15 seconds after ablation.

### 3D simulations

The simulation package used for 3D models was developed in Matlab and was adapted from Pierre et al. and Ershov et al.^14,44^. This package includes a module to extract and reconstitute 3D shapes, centrosome positions and polarity domains from segmented experimental imaging stacks, and to add predefined hypothesis for how microtubules pull in the cell. The distance between centrosomes, and the size of apical polar domains were defined directly from each experimental stack. Once parameters were defined as inputs, the model placed the centrosome pair in a random position and orientation, traced MTs from centrosomes to measure their length and associated a force to each, and computed both global forces and torques exerted by both MT asters. To search for minima, the simulation was based on a random walk, which follows the minimization of torques and forces. This was achieved by randomly modulating one of the 5 spatial parameters (3 for the position and 2 for the orientation angles), and recalculating the force torque at the subsequent step. The simulations followed the direction of force/torque minimization and stopped at a position/orientation once the iteration returns to this equilibrium after a given number of iterations (e.g. 300 runs). The length of the simulations, the duration needed to identify a stable equilibrium, the noise added to explore other parameters, and other intrinsic parameters of the loop, can be modulated, but were fixed for all the simulations performed in this work. Finally, to ensure that the centrosome position/orientation identified did not correspond to a local minimum which could have been biased by the initial conditions, the simulations were run typically 3-4 times from another random starting position/orientation. Finally, once a position was found, the model recalculated the torque landscapes as a function of the 2 possible angles.

Two general classes of simulations were performed. The first one, the “geometrical model” assumes that all MTs pull with a force that scales to the cube of their length (Fig 3). The second one, the “polarity model” assumes that all MTs pull with a force that scales to the cube of the length of MTs and that MTs that contact the apical polar domain exert an additional pulling force scaled to the square of the length of MTs, with a parameter β that represents the competition between geometry and polarity (see main text) (Fig 4 and Fig S3).

## Quantification and statistical analysis

### Cell sphericity

To segment cells in 3D and quantify cell sphericity, Z-stacks confocal acquisitions were segmented using the semi-automatic tool “membrane-based cell segmentation” of the IMARIS 9.2 software, based on membrane signals (Membrane-RFP). A background noise subtraction filter was applied to all acquisition. Errors were corrected manually and cells were classified as interphasic or mitotic based on MTs (using EMTB-GFP) and Hoechst signals.

### Centrosomes orientation and position 3D tracking

Centrosomes orientation and position in 3D coordinates were obtained manually with Fiji, from fixed or live Z-confocal-stacks. Centrosomes were assumed to localize at the centre of MT asters where MTs emerge (Fig S1D). The coordinates of the center of the apical cell membrane was manually annotated based on the signal of F-actin at the apical membrane. Then, 3D cell contour masks obtained with the Fiji-plugin “DM3D-Deforming Mesh”, were obtained to generate the 3D geometry of each cell, and the center of mass of this geometry was calculated using home-made Matlab codes (Mathwork)^44^. Centrosome positions were normalized to the apico-basal length of the cell in order to compare them between stages.

### Cell Volumes

Cell volumes at all stages were calculated by considering cells as either prolate or oblate spheroids, based on their aspect ratio in the plane of imaging.

### Nuclei position

Nuclei were tracked from confocal timelapses based on different nuclei markers (NLS-GFP, H2B-GFP, H2B-RFP or Hoescht). To obtain the *in vivo* nuclei distance from the cell surface we drew with Fiji a vector from the nuclei center to the center of the apical cell membrane. The nuclei-apical distance was measured at each time point to avoid errors due to cell shape changes and membrane contractions along the cell cycle. The net nuclear displacement in response to drug treatment or laser ablation was calculated as the difference between the nuclear-apical membrane distance at time 0s and that at the onset of metaphase. All distances were normalized to the AB cell length.

### Cytoplasm flows

To track cytoplasm flows, we labelled yolk granules which are dense in early embryos. For this, embryos were incubated in Nile blue at a 1/1000 dilution in ASW.F from fertilization, and imaged at the morula/blastula stage with spinning disk microscopy, following treatments with either DMSO or Nocodazole. Movies were analysed using the particle image velocimetry PIVlab tool in Matlab (Mathworks). The exterior of the cell was masked to be excluded from the analysis. Contrast limited adaptive histogram equalization and two-dimensional Wiener filter with accordingly windows of 20 and 3 pixels widths were applied on the images in the pre-processing steps for noise reduction. Image sequences were investigated in the Fourier space by four interrogation windows with 64, 32, 16, and 8 pixels widths and 50% overlapped area. The distribution of velocity components of vectors for each set was visually inspected and restricted to remove outliers in the post-processing stage. The output vector fields after smoothing were used for further analysis and for plotting flow maps and streamlines in Matlab.

### Statistical analysis

For all experiments, statistical analysis was performed using GraphPad Prism (GraphPad Software, Boston, Massachusetts USA). First, the normality of the data distribution (Gaussian distribution) was tested using a D’Agostino and Pearson test. When the data followed a normal distribution, we then compared them using parametric two-tailed unpaired Student’s t-tests (between two groups) or one one-way ANOVA and Tukey HSD post-hoc test (between >2 groups). When the data did not follow a normal distribution, we compared them using Mann–Whitney (2 groups) or Kruskal–Wallis (>2 groups) nonparametric tests. P<0.0001 (****), P<0.001 (***), P<0.01 (**); P<0.05 (*) and P ≥ 0.05 was considered not significant (ns). The number of embryos and experiments performed for all analysis are listed in the figure legends.

## KEY RESOURCES TABLE

| REAGENT or RESOURCE | SOURCE | IDENTIFIER |
| --- | --- | --- |
| **Antibodies** |  |  |
| YL1/2 rat monoclonal | Abcam | Ab6160 |
| DM1A mouse monoclonal | Abcam | Ab7291 |
| p-PKC $\zeta$ (Thr 410) rabbit polyclonal | Santa Cruz Biotechnology | sc-12894-R |
| Phospho-Myosin Light Chain2 (Ser 19) Rabbit polyclonal | Cell Signalling Technology | 3671 |
| Nano mouse Alexa Fluor™ 488 IgG1 | Thermo Fisher Scientific | SMS1AF488-1-100 |
| Goat anti-Rabbit IgG DyLight™ 488 | Invitrogen | 35552 |
| Donkey anti-Rat IgG Secondary Antibody, DyLight™ 650 | Invitrogen | SA5-10029 |
| **Bacterial and virus strains** |  |  |
| DH5α Competent Cells | Thermo Fisher Scientific | EC0112 |
| **Chemicals** |  |  |
| DMSO | Euromedex | UD8050 |
| Triton X-100 | Sigma-Aldrich | T8787 |
| HEPES | Sigma-Aldrich | H3375 |
| EGTA | Sigma-Aldrich | E3889 |
| Triton X-100 | Sigma-Aldrich | T8787 |
| Formaldehyde 16% (w/v), methanol free | Thermo Fisher Scientific | 28908 |
| NaBH<sub>4</sub> | Sigma-Aldrich | D282 |
| PABA | Sigma-Aldrich | A9878 |
| MgSO<sub>4</sub> | Sigma-Aldrich | M7506 |
| Nile Blue A | Sigma-Aldrich | N0766 |
| Glutaraldehyde Grade I, 25% | Sigma-Aldrich | G5882 |
| Glycerol | Sigma-Aldrich | G5516 |
| Protamine sulfate | Sigma-Aldrich | P4020 |
| Tween® 20 | Sigma-Aldrich | P1379 |
| CF 555/ Maleimide | Sigma-Aldrich | SCJ4600021 |
| Hoechst | Sigma-Aldrich | 33342 |
| Rhodamine Phalloidin | Invitrogen | R415 |
| Nocodazole | Sigma-Aldrich | M1404 |
| ML-7 | Sigma-Aldrich | I2764 |
| Latrunculin B | Sigma-Aldrich | 428020 |
| **Plasmids** |  |  |
| H2B-GFP | Lab Stocks |  |
| H2B-RFP | Lab Stocks |  |
| NLS-GFP | Lab Stocks |  |
| EMTB-3GFP | A gift from Alex McDougall (LBDV, Villefranche) |  |
| TMBD-mStaygold | This Study |  |
| Membrane-mCherry | Lab Stocks |  |
| Membrane-mScarlet3 | This Study |  |
| LifeAct-mScarlet3 | This Study |  |
| Others |  |  |
| Artificial seawater (Reef Crystal) | Instant Ocean | RC15-10 |
| Sigmacote | Sigma-Aldrich | SL2 |
| Software and algorithms |  |  |
| Image analysis software | FIJI |  |
| Statistical analysis | GraphPad Prism 10 |  |
| Matlab | MathWorks | mathworks.com/products/matlab.html; RRID:SCR_001622 |
| 3D models for centrosome positionning | Ershov and Minc<sup>44</sup> and Pierre et al.<sup>14</sup> | N/A |
| Cell segmentation | IMARIS 9.2 / DM3D Fiji Plugin |  |

