## Supplementary Material for "A Cell Size-Dependent Competition Between Geometry and Polarity Governs Nuclear and Spindle positioning in Early Embryos"

This supplementary material file contains 3 Supplemental figures, figure legends and supplemental Movie legends.

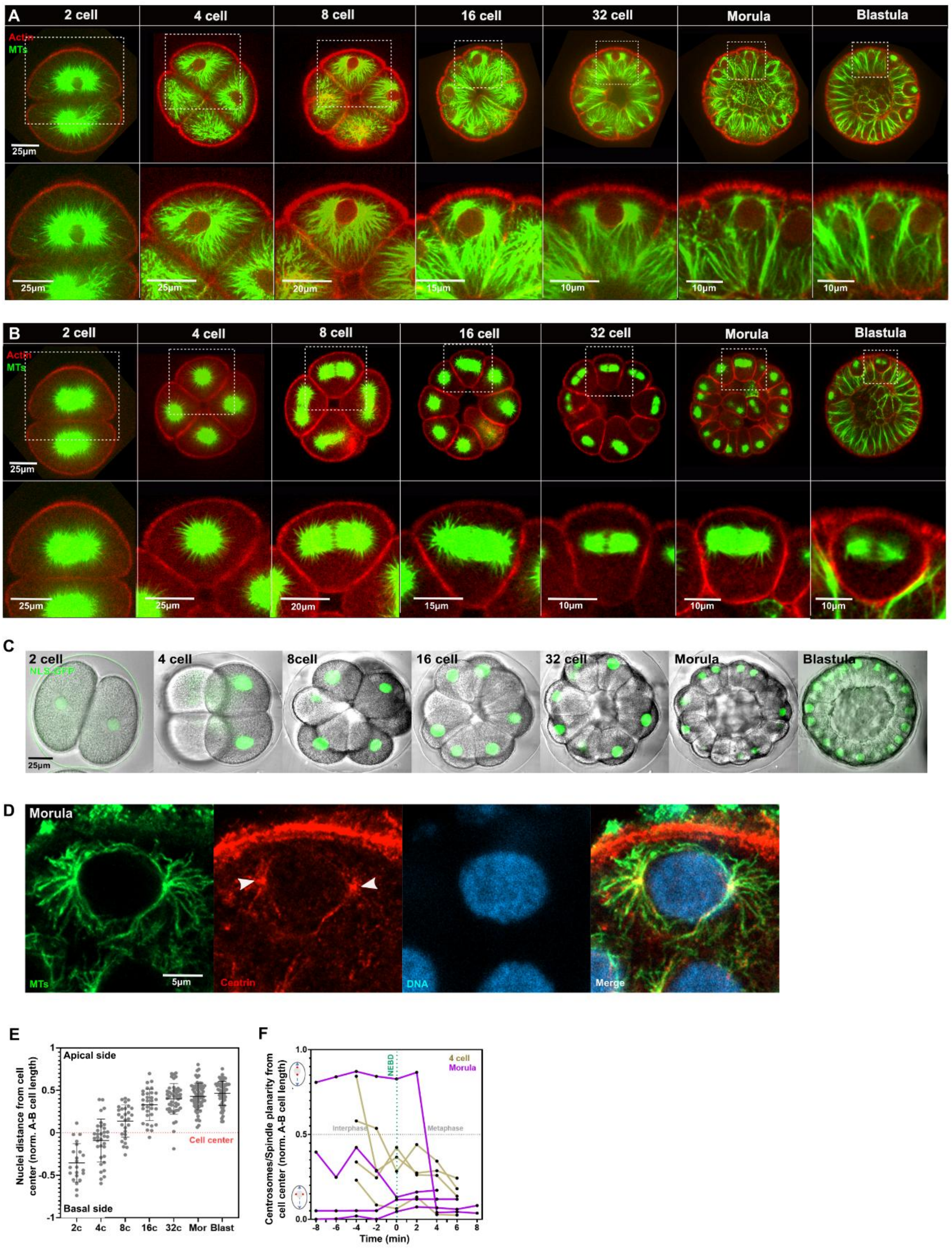

**Figure S1. Centrosome, nuclear and spindle positioning during early embryo development (Related to Figure 1). (A-B)** Time-lapse confocal acquisition of a sea urchin embryo injected with mRNA encoding for TMBD-StayGold and LifeAct-mScarlett to image MTs and F-actin respectively. Cells in interphase are depicted in A, and cells in metaphase in B. **(C)** Time-lapse confocal acquisition of a sea urchin embryo injected with NLS-GFP to visualize nuclei at subsequent developmental stages. **(D)** Confocal image of a fixed sea urchin blastomere at morula stage stained for MTs (green), Centrin (red) and DNA (blue). White arrows point at centrosomes. **(E)** Quantification of nuclear distances from the cell center (red dotted line), normalized to the apico-basal cell length across developmental stages. (n>15 cells/stage). **(F)** Single cell time evolution of interphase centrosome and spindle positions from interphase to metaphase at 4-cell and morula stages (n=4 cells/stage). Scale bar lengths are indicated in corresponding panels. Error bars represent +/- S.D.

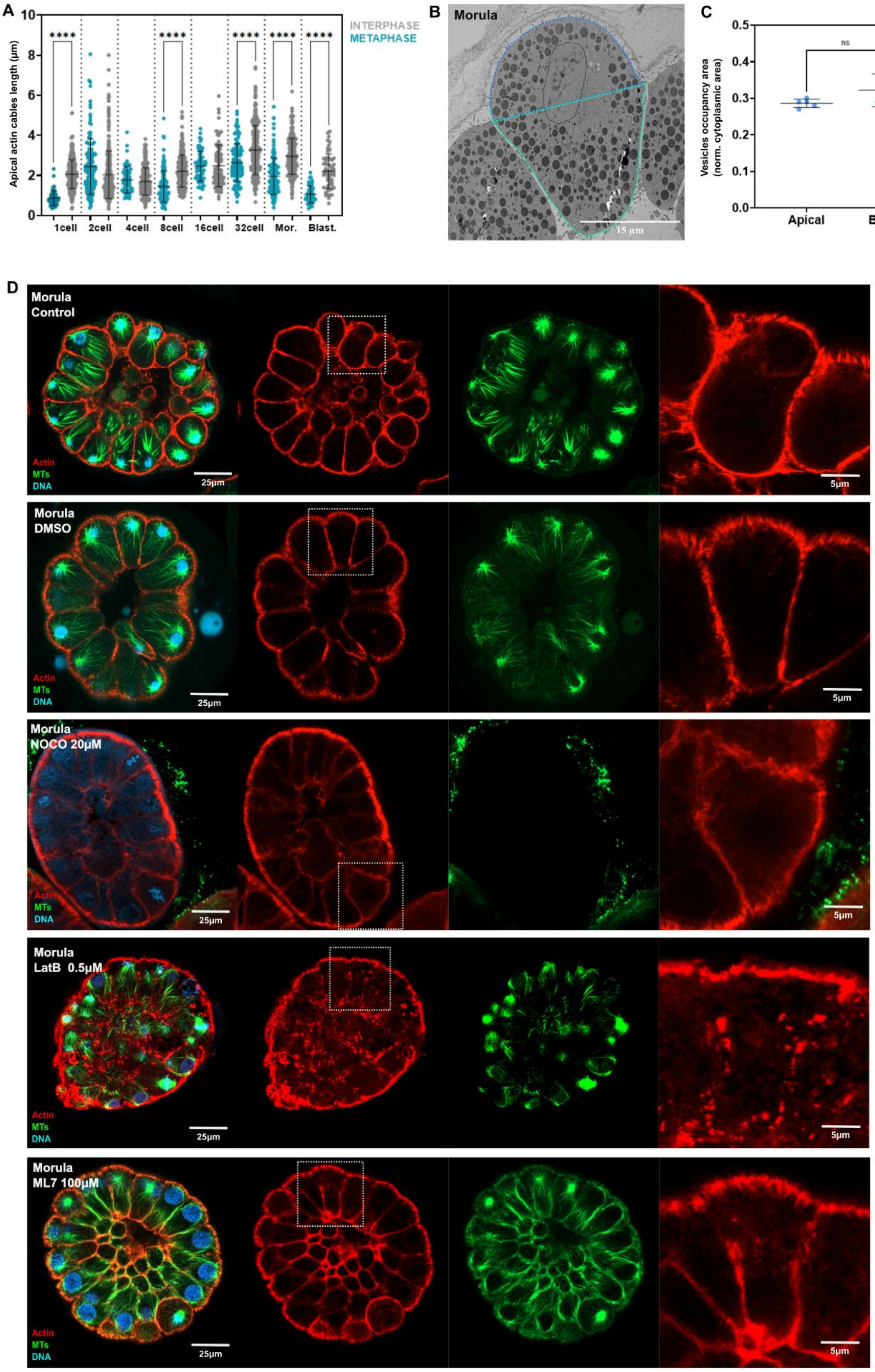
A
B
C
D
INTERPHASE
METAPHASE
Apical actin cables length (µm)
****
1cell
2cell
4cell
8cell
16cell
32cell
Mor.
Blast.
Morula
15 µm
ns
Vesicles occupancy area (norm. cytoplasmic area)
Apical
Basal
Morula Control
Morula DMSO
Morula NOCO 20µM
Morula LatB 0.5µM
Morula ML7 100µM
Actin
MTs
DNA
25µm
5µm

**Figure S2. Cytoskeleton and cytoplasm organization in developing embryos (Related to Figure 2). (A)** Apical actin cable length at different developmental stage in interphase/metaphase. **(B)** Transmission Electron Microscopy image of a blastomere of sea urchin morula. Blue and green dashed contours delineate the "apical" and "basal" cell areas considered for the quantification in (C). The black line delineates the nucleus**. (C)** Quantification of large organelles/vesicles crowding based on the total vesicle area normalized by the total apical/basal cytoplasmic cell area (n=5 cells). **(D)** Confocal acquisitions of fixed sea urchin embryos at morula stage treated by the indicated cytoskeletal inhibitors and stained for MTs (green) and F-actin (F-actin, red). White insets highlight the F-actin signal in individual blastomeres. Scale bar lengths are indicated in corresponding panels. Error bars represent +/- S.D. Results were compared using a two-tailed Mann–Whitney test. P-values are indicated as n.s., $P > 0.05$, ****, $P < 0.0001$.

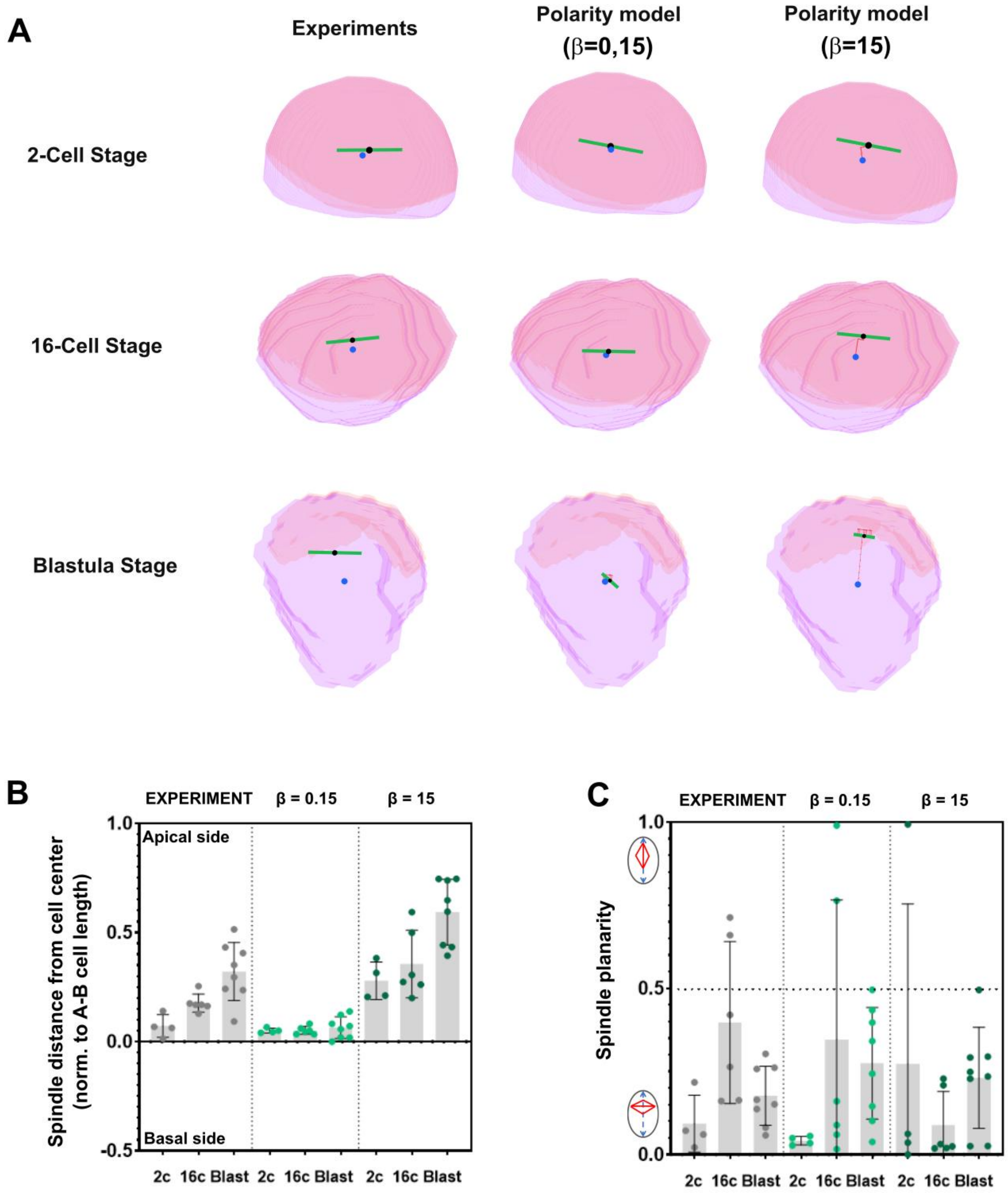
A
Experiments
Polarity model (β=0,15)
Polarity model (β=15)
2-Cell Stage
16-Cell Stage
Blastula Stage
B
EXPERIMENT
β = 0.15
β = 15
Spindle distance from cell center (norm. to A-B cell length)
Apical side
Basal side
1.0
0.5
0.0
-0.5
2c 16c Blast 2c 16c Blast 2c 16c Blast
C
EXPERIMENT
β = 0.15
β = 15
Spindle planarity
1.0
0.5
0.0
2c 16c Blast 2c 16c Blast 2c 16c Blast

**Figure S3 Influence of model parameters on centrosome positioning predictions (Related to Figure 4). (A)** 3D model predictions of metaphase spindle orientation and position at different stages, for low or high constant values of the parameter β. The green line marks the spindle axis, the blue dot, the cell center of mass and the black dot the spindle center, the pink area delineates the apical polar domain. **(B-C)** Experimental results (grey dots) and 3D model predictions for the same cells (light and dark green dots) for low or high constant values of the parameter β of spindle distance from the cell center normalized to apico-basal cell length (B) and spindle planarity (C) (n=4-8 cells/stage).

### Supplemental Movie Legends

**Movie S1 (Related to Figure 1)**: Time-lapse confocal projection of a sea urchin embryo injected with mRNA encoding for TMBD-StayGold and LifeAct-mScarlett to image MTs and F-actin respectively.

**Movie S2 (Related to Figure 1)**: Time-lapse confocal projection of a sea urchin embryo injected with NLS-mCherry to visualize nuclei at subsequent developmental stages. The fluorescent signal has been superimposed with DIC channel.

**Movie S3 (Related to Figure 2):** Time-lapse of embryos stained with Hoechst to visualize nuclei and treated at time 0 with nocodazole.
